# Exposure Duration Shapes the Hepatic Response to GenX: Divergent Acute and Chronic Transcriptomic Profiles Reveal Non-Monotonic Dose Effects and Increased Sensitivity in Human Liver Spheroids

**DOI:** 10.64898/2026.08.08.743693

**Authors:** Chanhee Kim, Abderrahmane Tagmount, Zhaohan Zhu, W. Brad Barbazuk, Rhonda Bacher, Christopher D. Vulpe

## Abstract

Hexafluoropropylene oxide dimer acid (GenX), a replacement for legacy per- and polyfluoroalkyl substances (PFAS), is increasingly detected in the environment, yet its chronic toxicity remains poorly characterized. Current safety assessments rely largely on short-term, high-dose studies that may not capture the biological consequences of long-term, low-dose exposure. To address this gap, we employed 3D human liver (HepG2/C3A) spheroids cultured in a continuously rotating bioreactor system (ClinoStar) to systematically evaluate dose- and time-dependent mRNA changes in response to GenX under environmentally relevant conditions. Spheroids were exposed to GenX (0.08-50 μM, spanning environmentally relevant to mechanistically informative concentrations) for acute (4 days) and chronic (4 weeks) durations, followed by genome-wide TempO-Seq transcriptomic profiling and benchmark dose (BMD) modeling. GenX elicited pronounced non-monotonic mRNA changes in acute exposure conditions, with the greatest number of differentially expressed genes (DEGs) observed at an intermediate concentration (0.4 μM). In contrast, chronic exposure exhibited a generally concentration-dependent increase in DEGs, except for the 10 μM condition, indicating a more consistent dose–response relationship than acute exposure. Notably, acute and chronic exposures elicited qualitatively distinct mRNA changes with low concordance across matched concentrations, demonstrating that exposure duration was a major determinant of mRNA changes. Acute low-dose GenX exposure preferentially modulated mRNA encoding components of cell cycle-related pathways, whereas acute higher dose exposures suppress mRNA levels of the constituents of lipid metabolic pathways and increase expression of mRNA encoding proteins involved in stress- and toxicity-associated signaling. Chronic exposure revealed a different pattern of changes in mRNA expression not observed under acute exposure conditions, including suppression of cellular components involved in lipid-related pathways at the lowest concentration tested. At higher concentrations, mRNA levels of components of multiple metabolic pathways were altered. Benchmark dose modeling identified a significantly lower transcriptomic point of departure (tPOD) for chronic exposure as compared to acute exposure, suggesting increased cellular sensitivity to prolonged GenX exposure and supporting the relevance of chronic models for human exposure assessment. Collectively, these findings demonstrate that GenX elicits time-dependent and non-monotonic changes in mRNA levels of human liver (HepG2/C3A) spheroids, with distinct responses depending on the exposure duration and dose. This study, therefore, highlights the importance of incorporating chronic, human-relevant *in vitro* models and transcriptomic endpoints into PFAS risk assessment and suggests that conventional short-term assays may underestimate the biological impact of sustained low-dose exposure.

**Key message (Impact of the study):** This study provides systematic comparisons of short term (4 day) versus longer term (4 weeks), environmentally relevant GenX exposure in human liver spheroids, revealing non-monotonic, time-dependent changes in mRNA levels encoding cellular components of lipid metabolism-related pathways with potential implications for appropriate dose and time exposure parameters for use in New Approach Methods to be applied in risk assessment.

## 1. Introduction

Perfluoroalkyl substances (PFAS) alternatives have been developed to replace ‘legacy’ PFAS that exhibit extreme environmental persistence and bioaccumulation in humans and wildlife due to their resistance to degradation^1^. Among these alternatives, hexafluoropropylene oxide dimer acid (HFPO-DA; GenX) was introduced as a replacement for perfluorooctanoic acid (PFOA) following increasing regulatory restrictions on legacy PFAS^2,3^. However, growing evidence indicates that GenX is also an emerging environmental contaminant of concern^4^ ^5^. Reflecting these concerns, the U.S. Environmental Protection Agency (EPA) released a human health toxicity assessment for GenX in 2021 and subsequently included GenX among PFAS subject to regulatory monitoring and drinking water standards^6^. More recently, the U.S. EPA announced the final drinking water regulation for six PFAS, including GenX, pointing out urgent human health concern of this chemical^7^. GenX has been detected in drinking water, groundwater, surface water, soil, and vegetation near fluorochemical manufacturing facilities, with concentrations in some contaminated regions exceeding recommended safety limites^8,9^. Despite increasing environmental detection, evidence of human exposure, and growing regulatory attention, the long-term health effects and human-relevant mechanisms of GenX toxicity remain insufficiently characterized^3,10,11^.

Accumulating epidemiological and experimental studies have identified potential adverse effects of GenX^12,13^, reporting GenX-induced toxicity in liver^14,15^, reproductive system^16,17^, endocrine systems^18^, development^19^, and immune function^20^. The liver is the primary site of GenX accumulation and one of its most sensitive toxicity targets^21,22^. Hepatic effects associated with GenX exposure include metabolic disruption, liver enlargement (i.e., hepatomegaly), altered lipid homeostasis, and extensive transcriptional reprogramming^23^. Transcriptomic and systems toxicology studies in rodents, fish, and in vitro models have demonstrated that GenX alters pathways involved in lipid metabolism, bile acid homeostasis, immune signaling, and cellular stress responses^24^ ^25–28^. Further, a recent study reported that GenX exposure suppress cholesterol biosynthesis and innate immune signaling in human liver spheroids^29^. Dysregulation of bile acid metabolism has also emerged as a potential contributor to GenX-induced hepatotoxicity, with *in vivo* studies demonstrating increased total bile acids, elevated primary bile acids, and reduced secondary bile acids following exposure^30,31^. Collectively, these findings indicate that GenX toxicity likely involves multiple interacting pathways and that a unified human-relevant mechanistic framework remains to be developed.

Importantly, most current knowledge of GenX toxicity is derived from acute or relatively short-term exposure studies conducted at moderate to high concentrations, particularly in case of *in vitro* approaches^2,32^. Such approaches may not adequately capture the biological consequences of prolonged low-dose exposure that more closely reflect real-world human exposure scenarios. Another limitation of previous studies is the lack of physiologically relevant human models capable of maintaining long-term hepatic function while capturing adaptive responses to chronic chemical exposure. Conventional two-dimensional (2D) hepatocyte cultures often fail to recapitulate the cellular architecture, metabolic competence, and long-term functionality of human liver tissue^33–35^. In contrast, three-dimensional (3D) human liver models, including HepG2 and HepaRG cell lines-derived or primary hepatocytes-derived spheroids, iPSC-derived spheroids/organoids provide a more predictive platform for chronic toxicity assessment by maintaining hepatic phenotype, supporting prolonged culture, and enabling more human-relevant toxicology evaluation^33,36^. Incorporation of environmentally relevant exposure concentrations further enhances the utility of these systems for human health risk assessment.

Most experimental studies investigating GenX toxicity, particularly those using human *in vitro* models, relied on relatively short exposure periods ranging from several hours to only a few days. For example, recent mechanistic studies examining GenX toxicity in human HepG2 cells and rat thyroid FRTL-5 cells employed exposure durations of up to 72 h^32,37^. Similarly, a recent transcriptomic study evaluating 24 different PFAS, including GenX, in human primary hepatocyte spheroids examined responses after 24 h and 10 days of exposure^29^. While these studies have provided valuable insights into the early molecular events and mechanisms underlying GenX toxicity, it remains unclear whether such relatively short-term exposure paradigms adequately capture the biological responses associated with sustained or chronic exposure.

Although short term exposure studies are useful for identifying early molecular events and mechanisms of action, their relevance to sustained human exposure remains uncertain because transcriptional responses may evolve through adaptation, cumulative perturbation, or transition to distinct biological states during chronic exposure. This issue is particularly important for quantitative transcriptomics, in which benchmark dose or concentration modeling is increasingly used to derive transcriptomic points of departure (tPODs) for chemical hazard and risk assessment. Short-term tPODs have shown encouraging concordance with chronic apical effects, which allowed it to be developed as efficient tools for screening and regulatory assessment^38^. However, tPOD estimates can vary with exposure duration in a chemical-dependent manner, and there remains no universal basis for assuming that a tPOD derived from an acute *in vitro* exposure will adequately represent the potency or biological processes associated with chronic exposure^39^. Direct comparison of acute and chronic exposure conditions is therefore needed to determine whether short-term transcriptomic responses provide a sufficiently protective and biologically representative basis for assessing chemicals such as GenX for which there is sustained human exposure.

In the present study, we systematically characterized dose- and time-dependent mRNA changes in response to GenX using a 3D human liver cell line (HepG2/C3A) spheroid model cultured in a rotating bioreactor system. To determine how exposure duration shapes biological responses, spheroids were exposed to GenX across environmentally relevant and mechanistically informative concentrations under acute (4 days) and chronic (4 weeks) exposure conditions. We hypothesized that prolonged exposure would alter both the magnitude and nature of GenX-induced mRNA changes, revealing biological effects that may not be predicted from conventional short-term studies. Through transcriptomic profiling and benchmark dose modeling, this study provides insight into the mechanisms and sensitivity of chronic GenX exposure and highlights the importance of exposure duration in the interpretation of transcriptomic data for PFAS risk assessment.

## 2. Methods

### 2.1. HepG2/C3A 3D spheroid culture and GenX exposure

HepG2/C3A cells (ATCC) were maintained in Minimum Essential Medium (MEM, Gibco) supplemented with 10 % fetal bovine serum (FBS), 1 % non-essential amino acids (Gibco), and 1 % Antibiotic-Antimycotic (Gibco) under standard culture conditions (37°C, 5% CO_2_). Three-dimensional (3D) spheroids were generated using the ClinoStar rotating bioreactor system (CelVivo) as previously described^33,36^. Briefly, HepG2/C3A cells were seeded onto a Spherical Plate 5D (SP5D) plate at 500 cells per microwell and allowed to aggregate for 24 h at 37°C and 5% CO_2_. The pre-aggregated spheroids were subsequently transferred into ClinoReactors and maintained under continuous rotation in the ClinoStar system. Culture medium was replaced two to three times per week throughout the experiment. To evaluate concentration- and duration-dependent transcriptomic responses, mature (a day after transfer of pre-aggregates) HepG2/C3A spheroids were exposed to hexafluoropropylene oxide dimer acid (HFPO-DA; GenX) at nominal concentrations of 0.08, 0.4, 2, 10, or 50 μM. Vehicle-treated (dimethyl sulfoxide, DMSO) spheroids served as controls. Two exposure durations were evaluated: acute exposure (4 days) and chronic exposure (4 weeks). For chronic studies, freshly prepared exposure medium containing GenX was replenished during routine medium changes throughout the exposure period. At the end of each exposure period, spheroids were harvested for transcriptomic analysis. Three independent biological replicates were collected for each treatment group.

### 2.2. TempO-Seq

Transcriptome profiling was performed using the TempO-Seq Human Whole Transcriptome Assay panel (BioSpyder Technologies Inc, Carlsbad, California) as previously described^36^. Briefly, total RNA was extracted and purified from each sample (spheroids) using a RNeasy Plus Mini Kit (Qiagen) following the manufacturer’s instructions. Throughout the RNA extraction procedure, we measured the 260/280 and 260/230 absorbance ratios to determine RNA purity and the presence of contaminants in our biological samples. Genomic DNA contamination was removed using DNase I (Ambion, UK) according to the manufacturer’s instructions. All sample O.D. values showed a 260/280 ratio of less than 1.8 and a 260/230 ratio of less than 2.0. The resultant RNA samples were then diluted with sterile water to obtain a consistent concentration of 100 ng/µL across all samples. An equal volume of 2X TempO-Seq Enhanced Lysis Buffer was added to each RNA sample, which provides an assay-ready sample in 1X TempO-Seq Enhanced Lysis Buffer. The assay-ready samples were annealed to the detector oligo (DO, targeting and binding a single RNA of interest) mix using the following steps: the assay-ready samples were incubated on plate at 70°C for 10 minutes, then ramped to 45°C for 50 minutes with the same ramp rate; the plate was maintained at 45°C for 16 hours, then cooled to 25°C. Annealing was followed by nuclease digestion (37°C for 1.5 hours, the type of nuclease is not indicated by BioSpyder protocol) to remove excess oligos, and ligating (by incubating for 1 hour at 37°C, then immediately raise to 80°C for 15 minutes) of a pair of DOs. The ligated product was PCR amplified using primers that contain sample tag (index) sequences and adaptors required for sequencing using the following. PCR cycles: 1. 37°C for 10 minutes, 2. 95°C for 1 minute, 3. 25 cycles of 95°C 10 sec; 65°C 30 sec; 68°C 30 sec, 4. 68°C for 60 sec, 5. Hold at 25°C. The amplified assay products are then pooled and purified/concentrated using the Macherey-Nagel NucleoSpin Gel and PCR Cleanup kit (Takara Bio USA, Inc, Mountain View, California). The library was sequenced using a NovaSeqX (performed at BioSpyder).

Primary data analysis was performed on a raw fastq files by BioSpyder using the TempO-SeqR (v.3.0) program. Initially, reads were extracted from the bcl files and demultiplexed with bcl2fastq v.2.20.0.42. The fastq files were processed with the TempO-SeqR (v.3.0), which uses star v.2.5 to align the reads, and the count function from QuasR to extract feature counts using a gtf reference. A raw gene_count table that details the signal levels (counts) for each probe in each sample was provided by BioSpyder.

### 2.3. Bioinformatics

All bioinformatics analyses for the raw gene_count data were conducted using integrated Differential Expression and Pathway analysis version 1.13 (iDEPv1.13)^40^. Counts were count per million (CPM) normalized and genes without 0.5 CPM in at least one sample were removed from analysis. To facilitate clustering analyses and PCA, the filtered gene count data were log2 transformed using the log_2_(cpm+c) function in the EdgeR package implemented in iDEPv1.13^40^. To assess sample variability, Pearson correlations of the log_2_(cpm+c) values were plotted to visually assess within sample similarity and between sample variability.

#### Differentially expressed genes

iDEPv1.13 was used to perform differential gene expression analysis on all pairs of sample groups using the DESeq2^41^ R package implemented in iDEPv1.13. Expression was filtered at a fold-change > 2 and a false discovery rate (FDR) threshold of < 0.05.

#### Biological pathway enrichment analysis

The resulting differentially expressed genes (DEGs) between comparison samples were then used to conduct enrichment analysis using Gene Ontology (Ensembl 92) and Kyoto Encyclopedia of Genes and Genomes (KEGG, Release 86.1) database. The background genes used for the enrichment analysis are the filtered genes from the genome-wide original gene list. Enrichment p-values are calculated based on a one-sided hypergeometric test, which is then adjusted for multiple testing using the Benjamini-Hochberg procedure and converted to FDR. The pathways are first filtered based on an FDR threshold of 0.05, and then the significant pathways are sorted by FDR. Functional analysis of DEGs was also conducted using STRING-DB_v12.0([STRING: functional protein association networks](https://string-db.org/cgi/input?sessionId=bf2AmUB4rACl&input_page_show_search=on), accessed 07/08/2026) for DEGs identified following 4-day and 4-week exposure to provide clustered networks. Upregulated and downregulated genes were analyzed separately. Gene symbols were mapped to *Homo sapiens* proteins, and only successfully mapped genes were included in the analysis. Functional enrichment was assessed using GO-BP annotations available in STRING, with the whole human genome used as the statistical background. Enriched GO-BP terms were ranked according to false discovery rate (FDR), and terms with an FDR < 0.05 were considered statistically significant.

#### Trendy analysis

Raw gene expression data was normalized using Median Normalization^42^ to account for variation in library size and sequencing depth across sample. Briefly, this method first calculates a ‘size factor’ for each sample accounting for variations in sequencing depth. Then, the method divides the expression values of each gene by the corresponding size factor. The result is a set of normalized expression values that are adjusted to a common scale across all samples.

To investigate transcriptomic responses across increasing GenX concentrations, we adapted the Trendy package^43^, which was originally developed for time-course gene expression analysis, to identify genes exhibiting non-linear dose-response relationships. In this framework, GenX concentration was treated as an ordered variable analogous to time. Trendy fits a series of segmented regression models to each gene and identifies statistically significant breakpoints representing changes in the direction or magnitude of expression responses across the dose range. Separate analyses were conducted for the acute (4 days) and chronic (4 weeks) spheroid exposure datasets. For each gene, the optimal segmented regression model was selected using the lowest Bayesian Information Criterion (BIC). Model fitting parameters included a maximum of three breakpoints, a minimum of five samples per segment, and a breakpoint significance threshold of p <0.2, consistent with the recommended Trendy settings. Genes identified by Trendy were interpreted as exhibiting distinct dose-responsive expression patterns, including monotonic, threshold-like, and non-monotonic responses.

### 2.4. Transcriptomic point-of-departure (tPOD) analysis

Transcriptomic point-of-departure (tPOD) analysis was performed using BMDExpress3.0^44,45^ ([BMDExpress - Sciome](https://www.sciome.com/bmdexpress/)) to quantify concentration-dependent transcriptional potency of GenX exposure in HepG2/C3A spheroids. Acute (4 days) and chronic (4 weeks) exposure datasets were analyzed independently to determine whether exposure duration alters transcriptomic sensitivity and pathway-level potency. The CPM-normalized gene expression data were imported into BMDExpress3.0 for benchmark concentration modeling. Genes were prefiltered using Williams trend test and fold-change criteria to identify transcripts exhibiting concentration-dependent responses. Genes passing the prefiltering step were modeled using Bayesian Model Averaging with Hill, Power, Exponential 3, and Exponential 5 models. A benchmark response corresponding to one standard deviation from the control mean was applied. Modeled genes were subsequently filtered to remove poorly fitted or unreliable benchmark concentration estimates, including genes with benchmark concentrations greater than the highest tested concentration and genes with excessive uncertainty based on BMCU (Upper)/BMCL (Lower) ratios^46^. Genes passing all pre-and post-filtering criteria were retained for downstream tPOD analysis. For each exposure duration, tPOD values were estimated using gene-level benchmark concentration distributions and pathway-level benchmark concentration analysis. Gene-level benchmark concentrations were used to derive the most sensitive tPOD. Pathway-level tPODs were calculated from biological pathways containing multiple responsive genes, and the lowest reliable pathway-level tPODs were interpreted as the lowest concentration associated with biological pathways perturbation. Acute and chronic tPODs were compared to evaluate whether chronic GenX exposure increased transcriptomic sensitivity (i.e., lower concentration to meet criteria for tPOD relative to short-term exposure).

## 3. Results

### 3.1. GenX-induced transcriptional responses are both dose- and time-dependent, with unexpected reversals in DEG magnitude across doses

Figure 1A summarizes the overall experimental approach used in this study. We used a ClinoStar bioreactor system (CelVivo) to continuously rotate HepG2/C3A spheroids generated by pre-aggregation procedure (see Methods), supporting prolonged cultures. The HepG2/C3A spheroids were then exposed to GenX, either acute (4 days) or chronic (4 weeks) from environmentally relevant (0.08 µM) to mechanistically informative concentrations (50 µM)^47,48^. Although 0.08 µM concentration exceeds typical levels detected in human serum^49,50^, it falls within the range of reported concentrations in water at contaminated sites^51,52^ and was previously used in *in vitro* studies to account for limited toxicokinetic processes^29,53^, enabling detection of early molecular perturbations. After exposures, RNAs from both dose-dependent acute and chronic samples were used for whole-transcriptome profiling via TempO-Seq followed by pathway enrichment and tPOD comparison analyses (Figure 1A).

**Figure 1.**
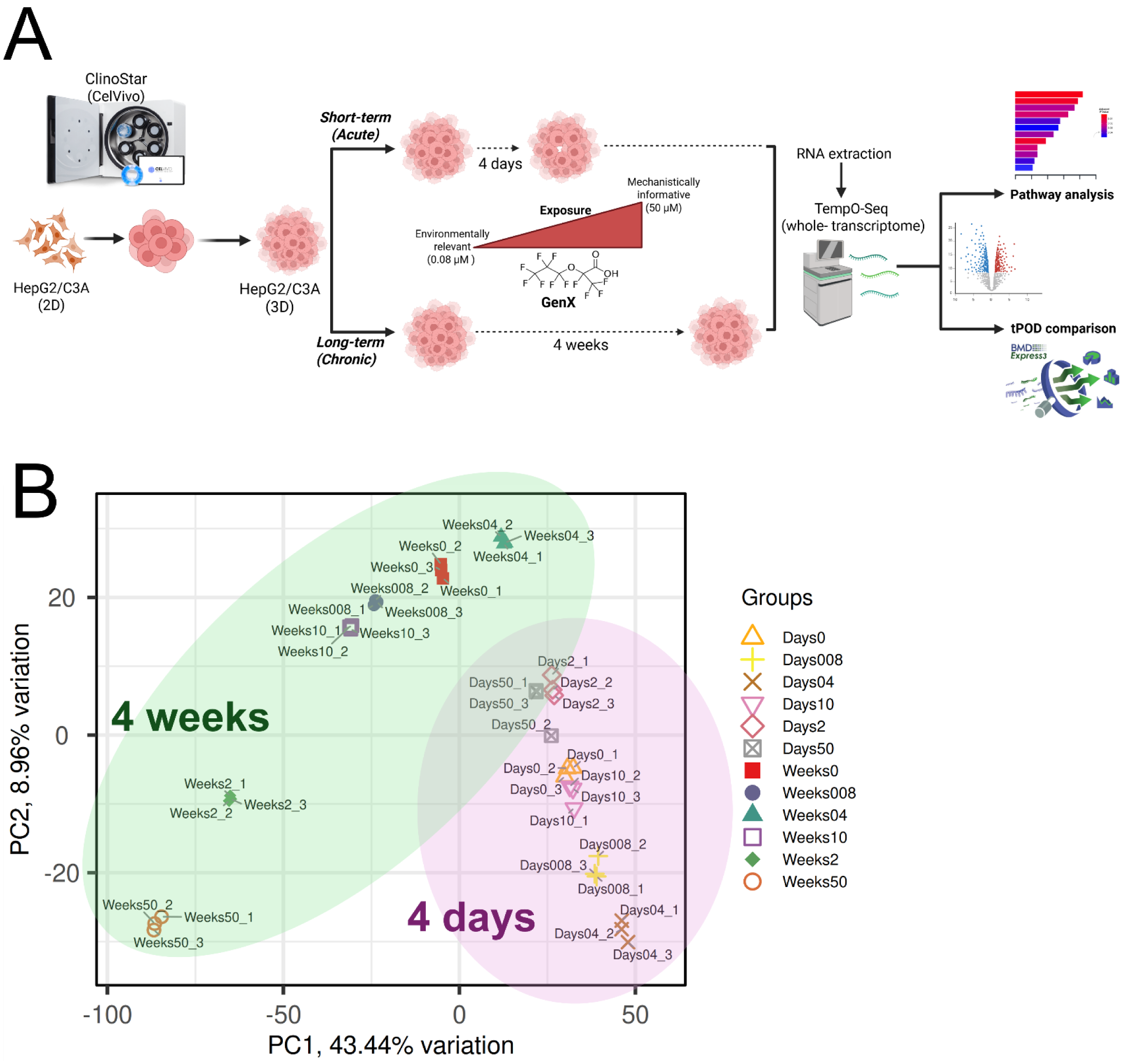
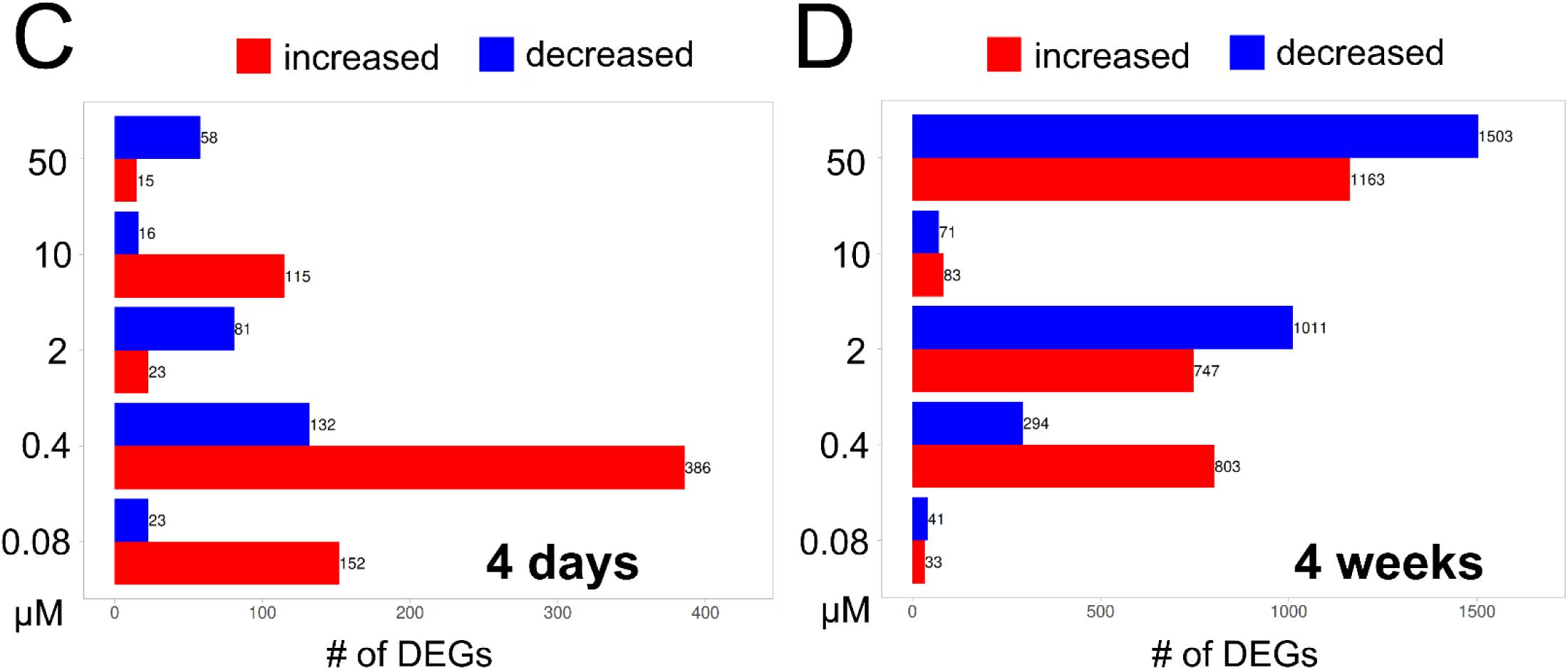
Experimental design & global transcriptional overview. (A) Schematic experimental design of this study. 3D spheroids generated using HepG2/C3A cells were exposed to GenX e for 4 days (acute) or 4 weeks (4 weeks) at different doses from an environmentally relevant dose (0.08 μM) to a mechanistically informative dose (50 μM). Collected samples were used for TempO-Seq (transcriptome profiling), followed by pathways-level enrichment and tPOD comparison analyses. (B) PCA plot of all samples analyzed in this study. (C)/(D) Comparison of differentially expressed genes (DEGs) in acute (4 days) and chronic (4 weeks) GenX exposed HepG2/C3A spheroids across doses (DEGs criteria: Log_2_FC>1 and FDR<0.05).

Dimension reduction of the data from all of spheroid RNA samples of both acute and chronic GenX exposure via PCA revealed that acute and chronic sets are clearly separated (Figure 1B). Notably, chronic exposure samples are more separated from each other than the acute counterparts (Figure 1B). HepG2/C3A 3D spheroids exposed to GenX for 4 days or 4 weeks exhibited varying numbers of differentially expressed genes (DEGs) across GenX concentrations (0.08 μM to 50 μM). For acute exposure to GenX, DEGs of 0.08 μM, 0.4 μM, 2 μM, 10 μM, 50 μM were 161 (Up: 142, Down: 19), 476 (Up: 349, Down: 127) 103 (Up: 15, Down: 88), 87 (Up: 71, Down: 16), and 65 (Up: 6, Down: 59), respectively (Figure 1C). The DEGs of acute GenX exposure in HepG2/C3A spheroids showed a non-monotonic dose-response (NMDR) (Figure 1C) which is defined as a dose-response curve in which the slope changes sign at least once over the range of doses examined^54,55^. For the chronic exposure scenario, the pattern of DEGs looked more monotonic except for the dose of 10 μM (Figure 1D). DEGs of 0.08 μM, 0.4 μM, 2 μM, 10 μM, 50 μM were 59 (Up: 32, Down: 27), 983 (Up: 789, Down: 294), 1711 (Up: 749, Down: 962), 157 (Up: 97, Down: 60), and 2656 (Up: 1168, Down: 1488), respectively (Figure 1D). Together, acute and chronic exposure to GenX shows qualitatively different transcriptomic responses, indicating NMDR behavior in human HepG2/C3A spheroids.

### 3.2. Conserved GenX-responsive genes exhibit distinct transcriptional dynamics during acute and chronic exposure

To identify changes in mRNA level consistently associated with GenX exposure regardless of concentration, we focused on genes that were differentially expressed (either increased or decreased mRNA levels) across all exposure concentrations within each exposure duration. These constitutively responsive genes were further analyzed using a modified application of Trendy analysis^43^ to characterize their expression dynamics across the concentration gradient (see Methods). We identified 8 constitutively increased GenX-responsive transcripts and 9 decreased transcripts in acute exposure (Figure 2), while 132 increased and 186 decreased transcripts in chronic exposure (Figure 3).

**Figure 2.**
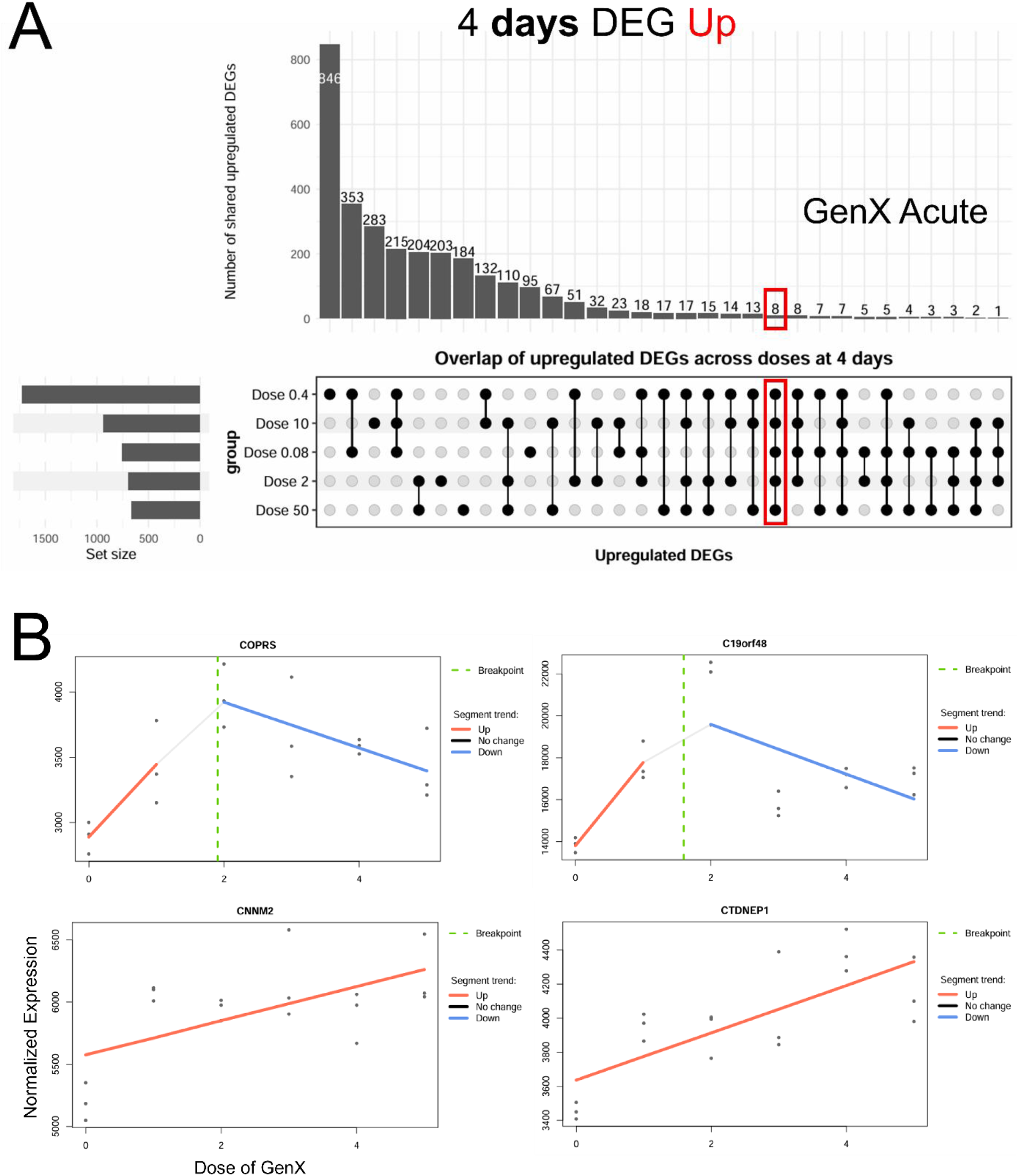
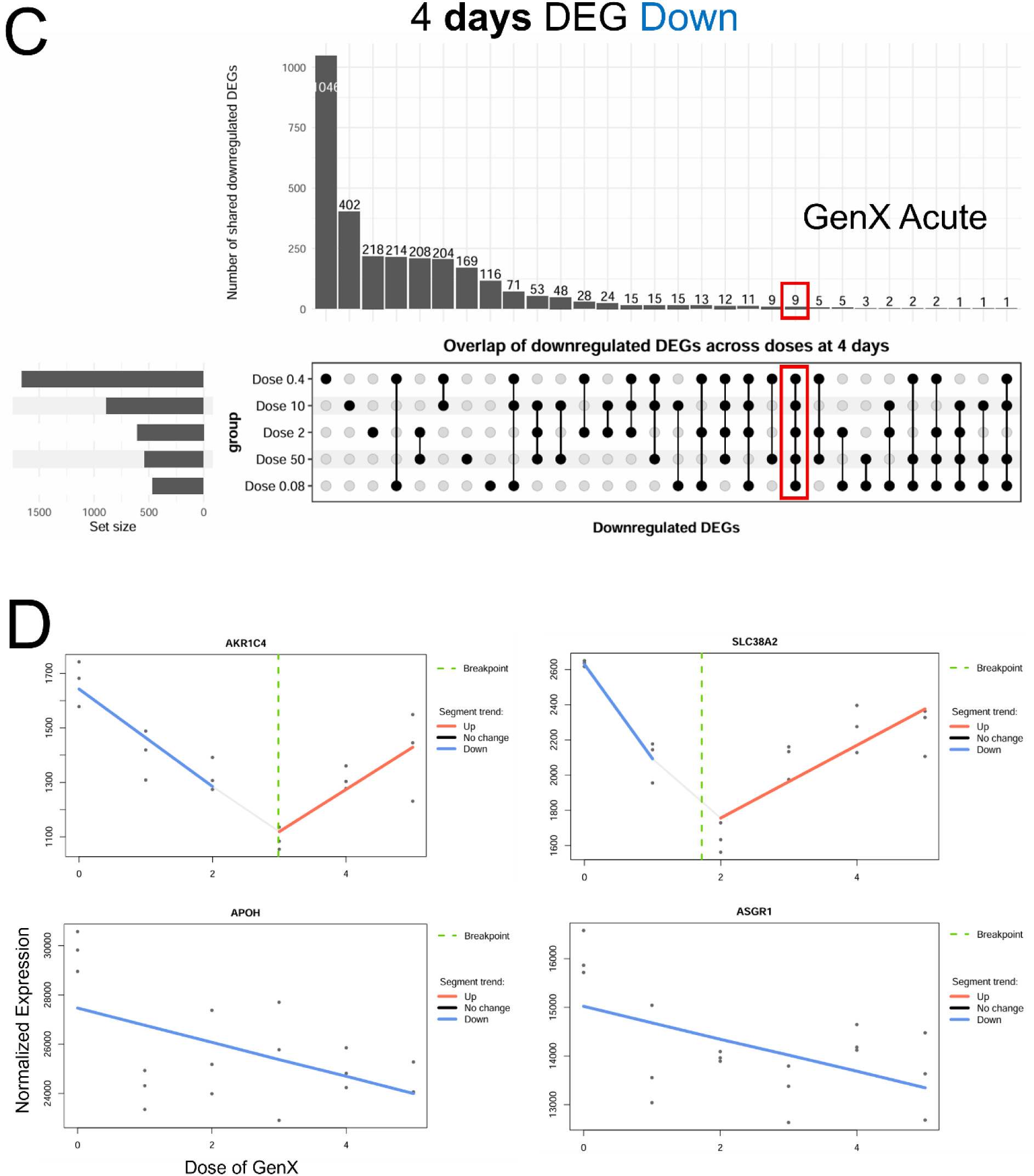
Conserved mRNA changes across GenX concentrations during acute (4 days) exposure condition. UpSet plots (A and C) showing the overlap of significantly altered genes (either increased (“Up-regulated”) or decreased mRNA expression (“Down-regulated”)) among HepG2/C3A spheroids exposed to GenX for 4 days at 0.08, 0.4, 2, 10, and 50 μM. Horizontal bars indicate the total number of differentially expressed genes (DEGs) (FDR <0.05) identified at each concentration, whereas vertical bars represent the number of genes shared among the indicated dose combinations. Constitutively altered transcripts across all GenX concentrations during acute exposure were subjected to Trendy segmented regression analysis^43^. Representative expression patterns illustrate transcripts exhibiting monotonic, threshold-like, and non-monotonic dose-dependent changes across the exposure range (B and D). Expression values are presented as normalized counts. Solid lines indicate fitted segmented regression models generated using Trendy.

**Figure 3.**
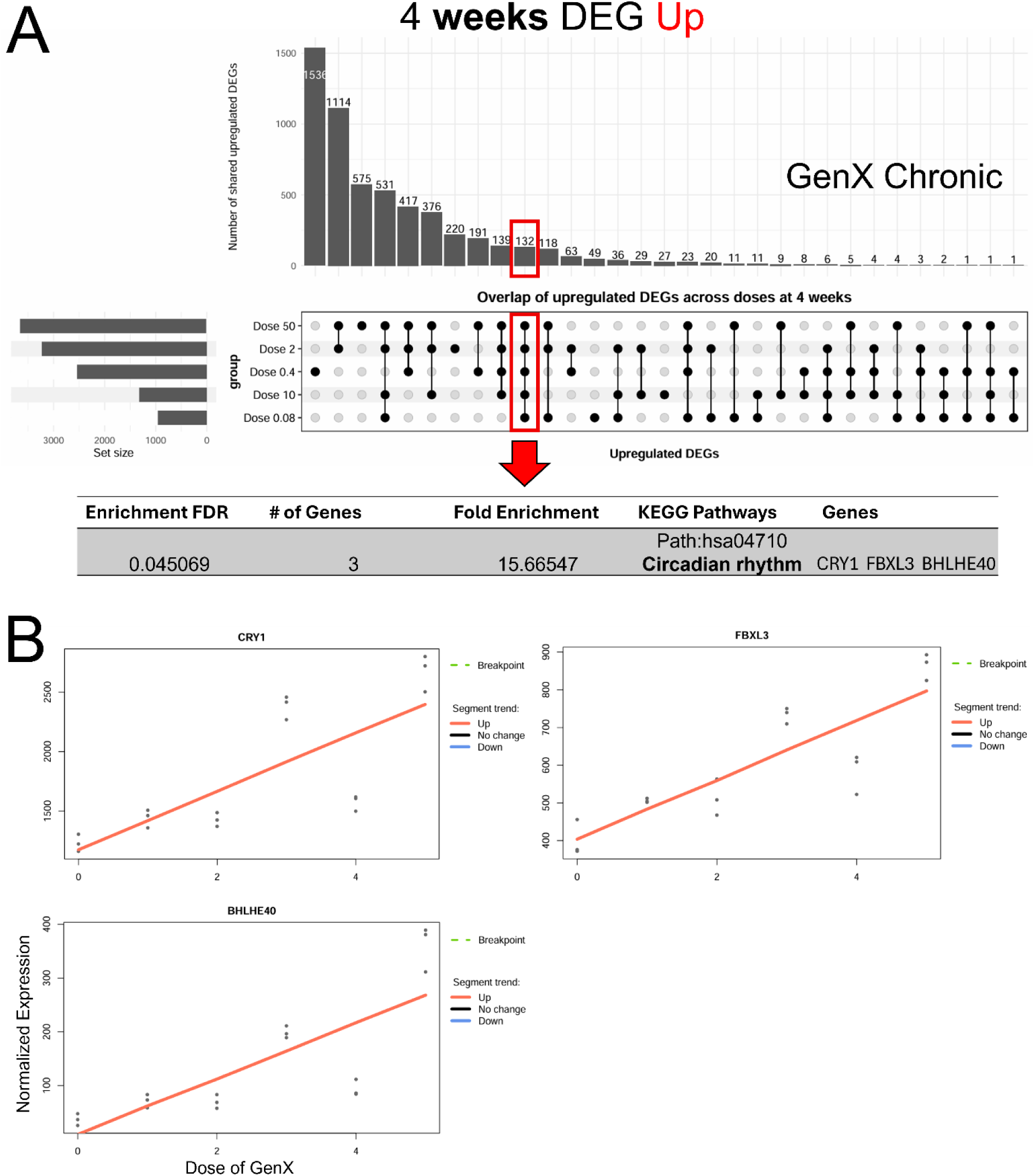
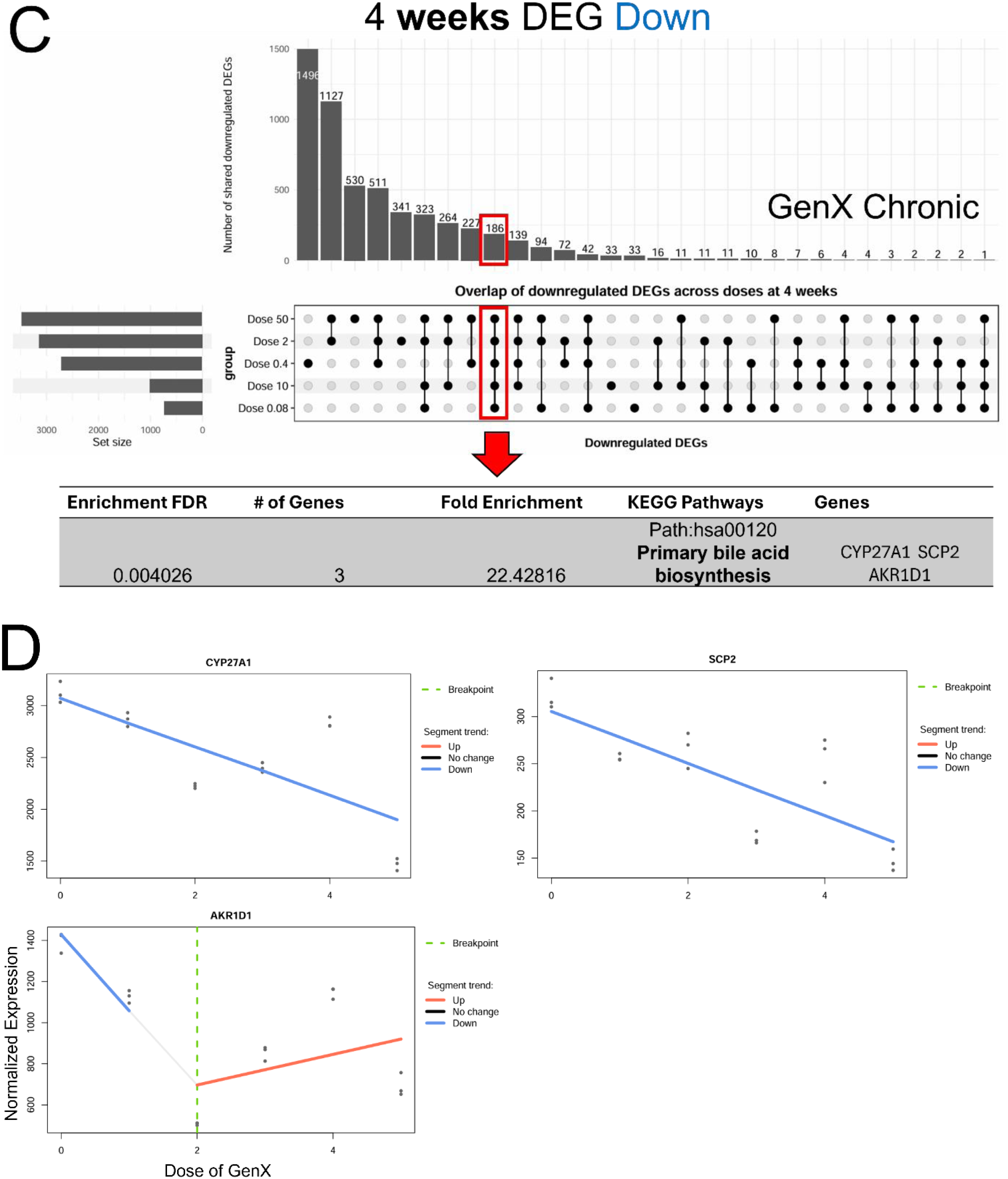
Conserved mRNA changes responses across GenX concentrations during chronic (4 weeks) exposure condition. UpSet plots (A and C) showing the overlap of significantly altered transcripts (either increased (UP DEGs) or decreased (DOWN DEGs) mRNA expression) among HepG2/C3A spheroids exposed to GenX for 4 weeks at 0.08, 0.4, 2, 10, and 50 μM. Horizontal bars indicate the total number of differentially expressed genes (DEGs) (FDR <0.05) identified at each concentration, whereas vertical bars represent the number of transcripts shared among the indicated dose combinations. Constitutively altered transcripts across all GenX concentrations during chronic exposure were subjected to Trendy segmented regression analysis^43^. Representative expression patterns illustrate transcripts exhibiting monotonic, threshold-like, and non-monotonic dose-dependent changes across the exposure range (B and D). Expression values are presented as normalized counts. Solid lines indicate fitted segmented regression models generated using Trendy.

Acute exposure revealed multiple classes of dose-responsive behavior, including monotonic, threshold-like, and non-monotonic expression patterns (Figure 2A). Several transcripts exhibited maximal responses at intermediate concentrations, consistent with the non-monotonic DEG distribution observed in Figure 1. Trendy analysis identified several transcripts exhibiting distinct dose-dependent mRNA level changes following acute GenX exposure. While most responsive transcripts displayed biphasic (i.e., adaptive) patterns with an initial increase followed by stabilization or decline (e.g., *COPRS, C19orf48*), a small subset of transcripts, including *CNNM2* and *CTDNEP1*, showed a sustained monotonic increase across the entire concentration range (Figure 2B). These findings suggest that acute GenX exposure elicits both adaptive transcriptional or other mRNA regulatory responses as well as a limited number of robust concentration-specific molecular responses. Analysis of the acute decreased transcripts (Figure 2C) identified a subset of transcripts exhibiting biphasic trajectories characterized by an initial decrease followed by partial recovery at higher concentrations, consistent with adaptive or non-monotonic regulatory responses, including *AKR1C4* and *SLC38A2* (Figure 2D). In contrast, several transcripts, such as *APOH* and *ASGR1,* displayed sustained concentration-dependent decreases across the entire GenX exposure range (Figure 2D).

Compared to acute exposure, chronic exposure generated more stable and sustained transcriptional trajectories, characterized by progressive increases or decreases across increasing concentrations (Figure 3). Of note, due to the substantially larger number of constitutive DEGs in the condition of chronic exposure, we first sought any enriched biological pathways in those DEGs. For increased DEGs, circadian rhythm was the top KEGG enriched pathway, allowing us to prioritize the member genes (*CRY1*, *FBXL3*, and *BHLHE40*) (Figure 3A and 3B). For decreased DEGs, primary bile acid biosynthesis was identified as a top KEGG enriched pathway, and we focused on member genes (*CYP27A1*, *SCP2*, and *AKR1D1*) for the Trendy analysis (Figure 3C and 3D). Trendy analysis of chronically increased transcripts revealed significant enrichment of circadian rhythm pathway (Figure 3A). Notably, core circadian regulators, including *CRY1, FBXL3*, and *BHLHE40,* exhibited progressive increases across the GenX concentration range (Figure 3B), suggesting that long-term GenX exposure induces a coordinated transcriptional or other mRNA regulatory program involving circadian clock regulation. These findings further support the notion that exposure duration fundamentally influences the molecular response landscape and may determine whether early adaptive responses transition into sustained biological reprogramming. Constitutively decreased transcripts across increasing GenX concentrations during chronic exposure condition identified a coordinated set of transcripts encoding cellular components involved in primary bile acid biosynthesis that exhibited dose-dependent mRNA level alterations (Figure 3C). Notably, *CYP27A1* and *SCP2* displayed progressive decreases in expression, whereas *AKR1D1* exhibited a biphasic response characterized by an initial decrease followed by partial recovery at higher concentrations (Figure 3D). Given the central roles of the corresponding gene products in cholesterol turnover, intracellular sterol transport, and bile acid biosynthesis^56^, these patterns collectively suggest that disruption of bile acid homeostasis represents a sensitive molecular response to chronic GenX exposure. Collectively, although concentration-specific mRNA changes dominated both exposure durations, chronic exposure produced substantially larger sets of transcripts shared across multiple concentrations, suggesting convergence toward stable mRNA changes during prolonged GenX exposure.

### 3.3. Acute Low-dose GenX exposure drives robust changes in gene expression in human hepatic spheroids

We next characterized concentration-dependent biological responses following acute GenX exposure based on KEGG and Gene Ontology Biological Process (GO-BP) enrichment analyses of gene products of differentially expressed genes (Figure 4). We inferred distinct biological changes across the concentration range, revealing a progressive transition from mRNA changes consistent with a cell proliferative response at low concentrations to more metabolic and signaling perturbations at higher concentrations. At environmentally relevant concentrations (0.08 and 0.4 µM), enriched pathways were dominated by cell cycle-related processes. KEGG analysis identified cell cycle-associated pathways among genes whose mRNA expression level increased, while GO-BP analysis showed significant enrichment of chromosome segregation, mitotic nuclear division, and cell cycle progression, indicating an early adaptive transcriptional response or other mRNA regulatory response (Figure 4A-B). Additional GO-BP enrichment analysis using STRING-DB at 0.08 µM further supported these findings, highlighting interconnected biological processes involved in mitotic cell cycle regulation, chromosome organization, and cell division, confirming that low-dose GenX primarily perturbs proliferative regulatory networks.

**Figure 4.**
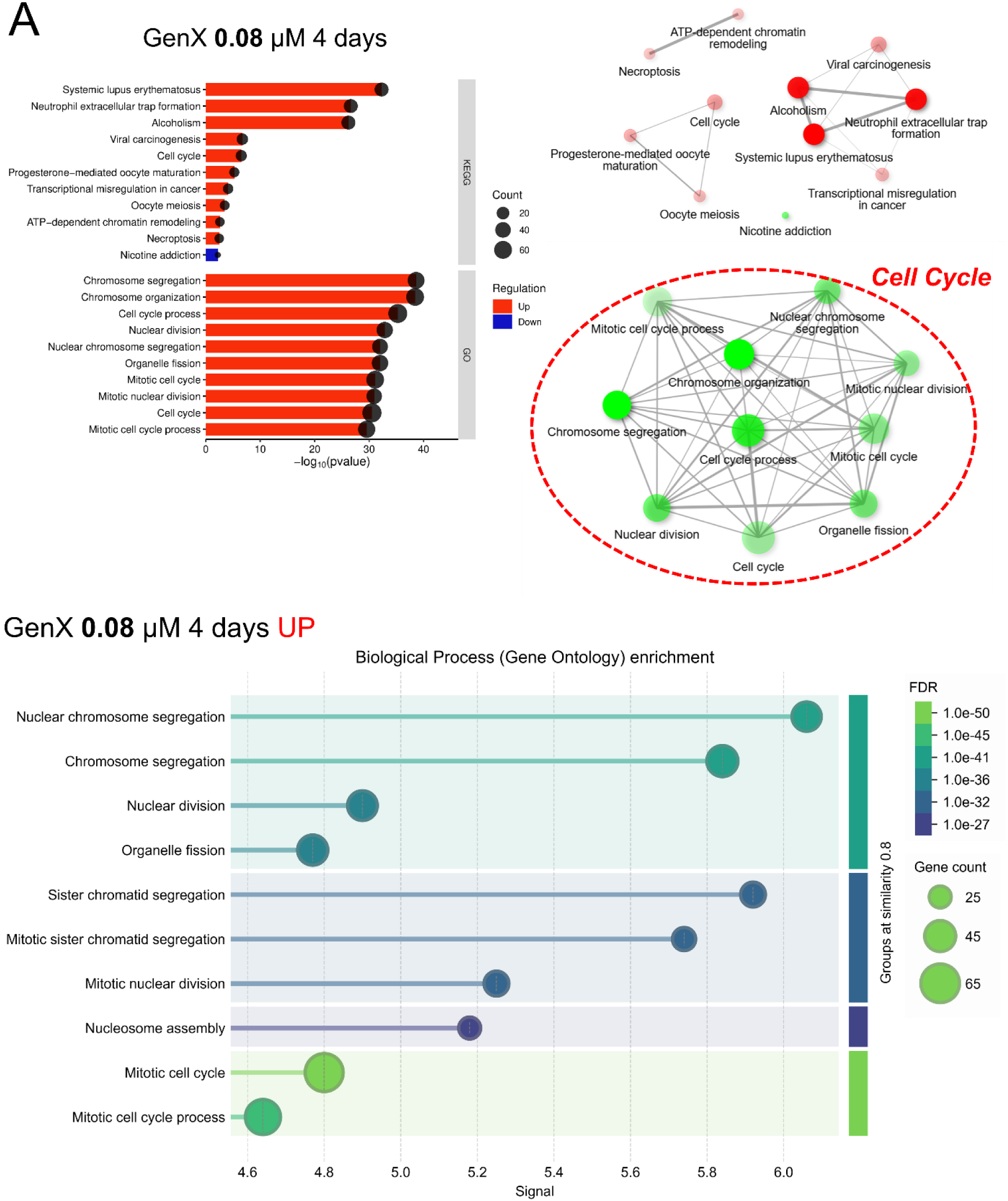
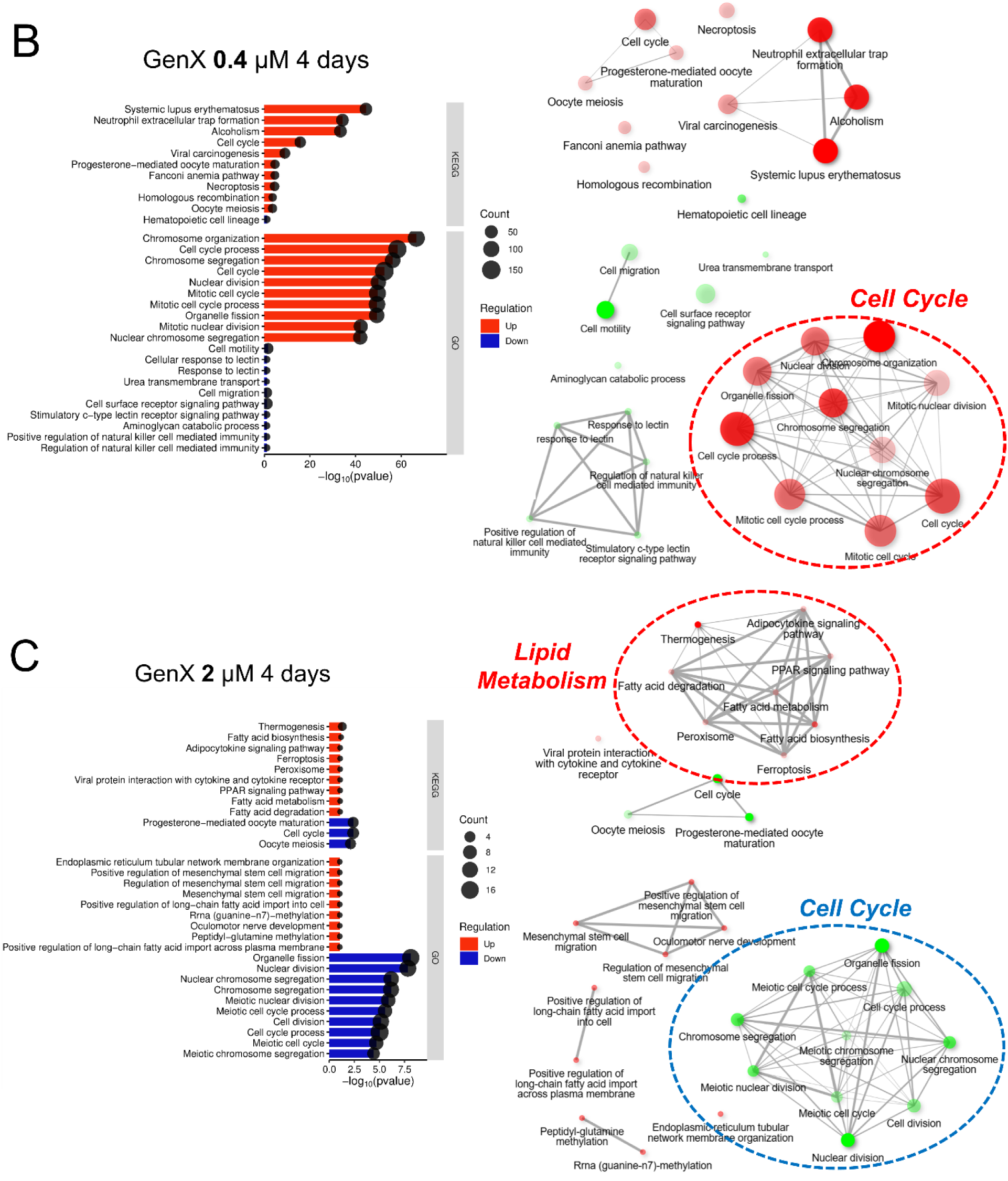
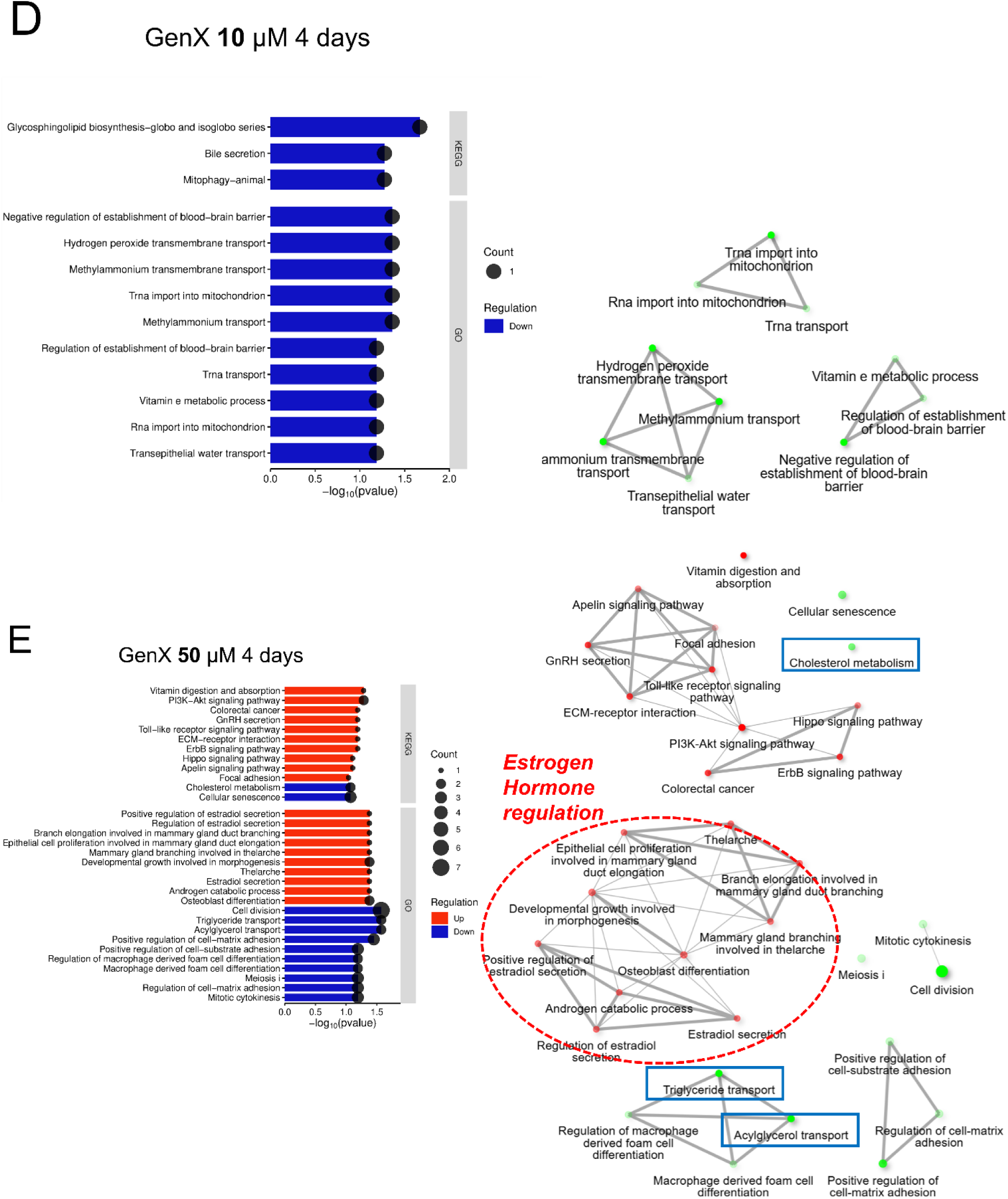
Acute GenX exposure induces concentration-dependent mRNA changes consistent with a transition from cell cycle regulation to lipid metabolism and endocrine-related pathways. Functional enrichment analysis of differentially expressed genes following 4 days of GenX exposure at 0.08 (A), 0.4 (B), 2 (C), 10 (D), and 50 (E) μM, respectively. Representative enriched KEGG and Gene Ontology Biological Process (GO-BP) terms are shown for each concentration with graphical presentation of clustered KEGG pathways and GO-BP. For the lowest concentration (0.08 μM), additional GO-BP enrichment analysis generated using STRING-DB is also presented. Enriched pathways were identified using DEGs (|log2FC| > 1, FDR < 0.05). Only the top 10 enriched pathways are shown.

At the intermediate concentration (2 µM), pathway enrichment of the protein products of the DEGs shifted toward lipid metabolism while maintaining enrichment of cell cycle-related processes, suggesting the onset of metabolic perturbation accompanying continued proliferative responses (Figure 4C). In contrast, relatively few significantly enriched pathways were detected at 10 µM despite measurable mRNA changes, consistent with a transient reduction in coordinated pathway-level responses (Figure 4D). At the highest concentration (50 µM), pathway enrichment shifted toward endocrine- and development-associated biological processes, including regulation of hormone levels and hormone metabolic processes, together with activation of PI3K-Akt, Toll-like receptor, and Hippo signaling pathways and suppression of cholesterol metabolism pathways (Figure 4E). Collectively, these findings demonstrate that acute GenX exposure induces concentration-dependent biological responses characterized by a progressive transition from early mRNA changes encoding proteins involved in cell cycle regulation at low concentrations to lipid metabolic perturbation and ultimately stress- and endocrine-associated signaling pathways at higher concentrations.

### 3.4. Chronic (4 weeks) GenX exposure reveals *qualitatively* distinct transcriptional programs

Functional enrichment analysis of proteins encoded by DEGs following 4 weeks of GenX exposure demonstrated a progressive concentration-dependent remodeling of biological processes that differed markedly from the acute response (Figure 5). In contrast to acute exposure, where low concentrations primarily affected cell cycle-associated pathways, chronic exposure initiated coordinated mRNA changes consistent with metabolic alterations even at the lowest concentration tested, indicating increased biological sensitivity to prolonged GenX exposure.

**Figure 5.**
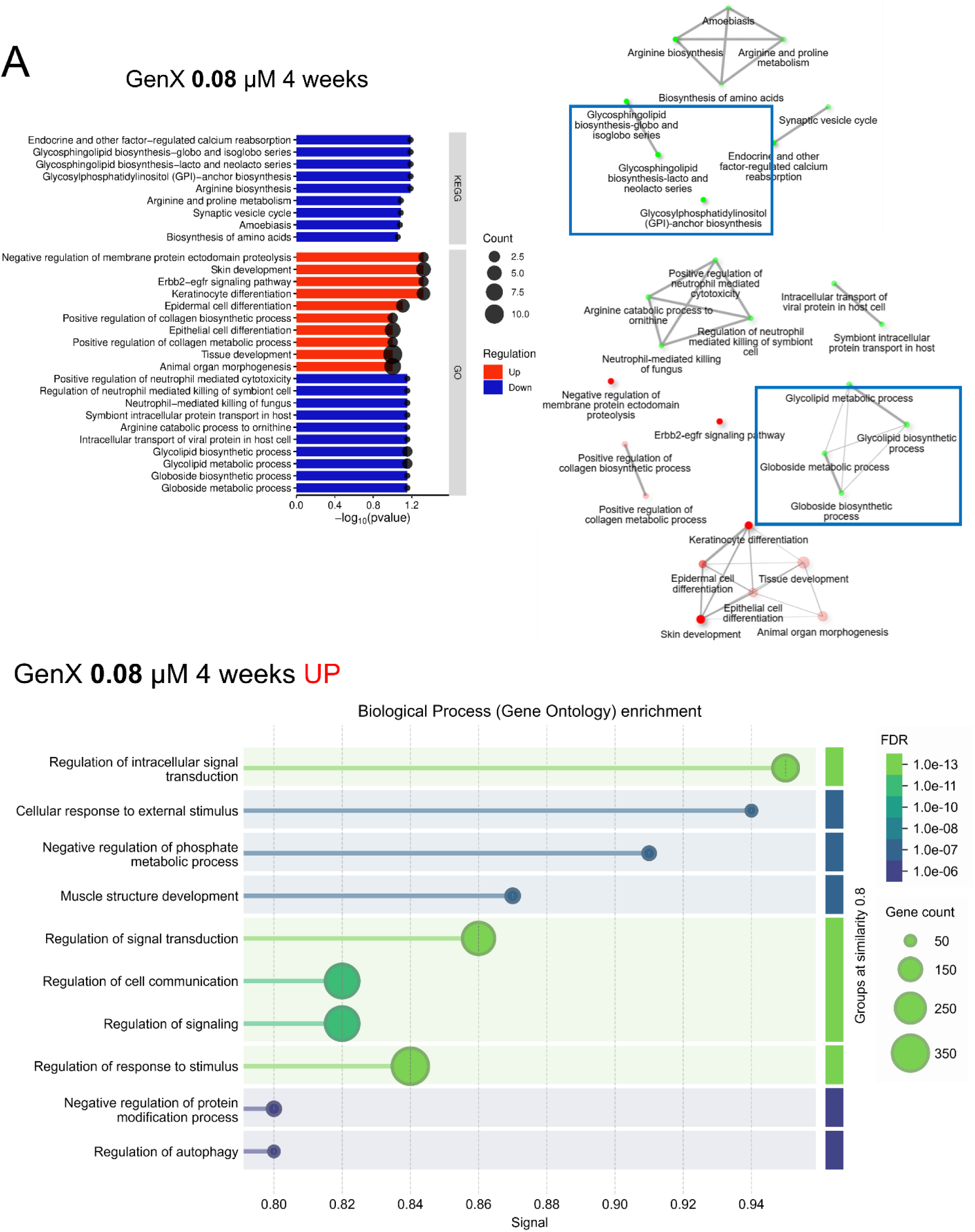
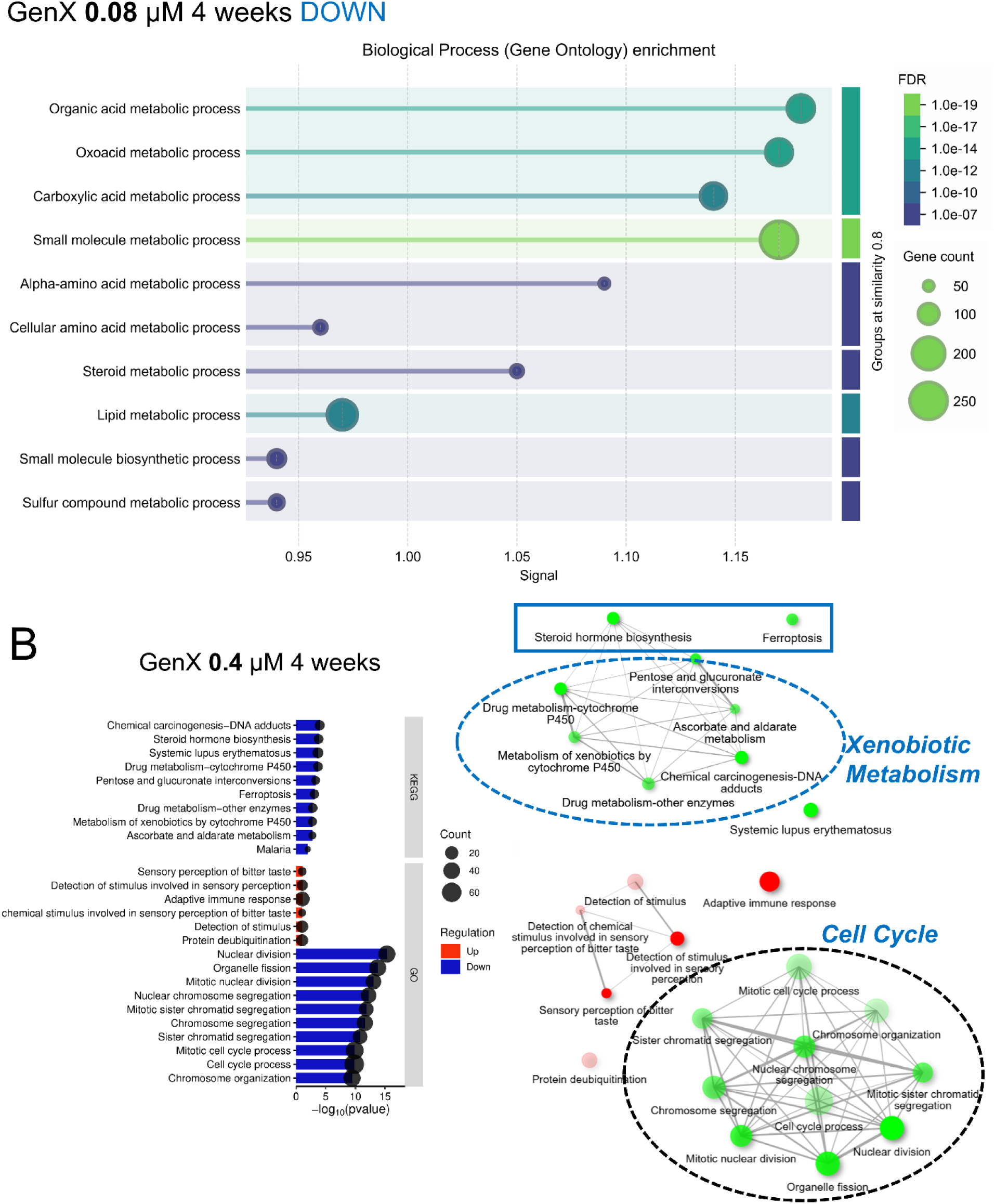
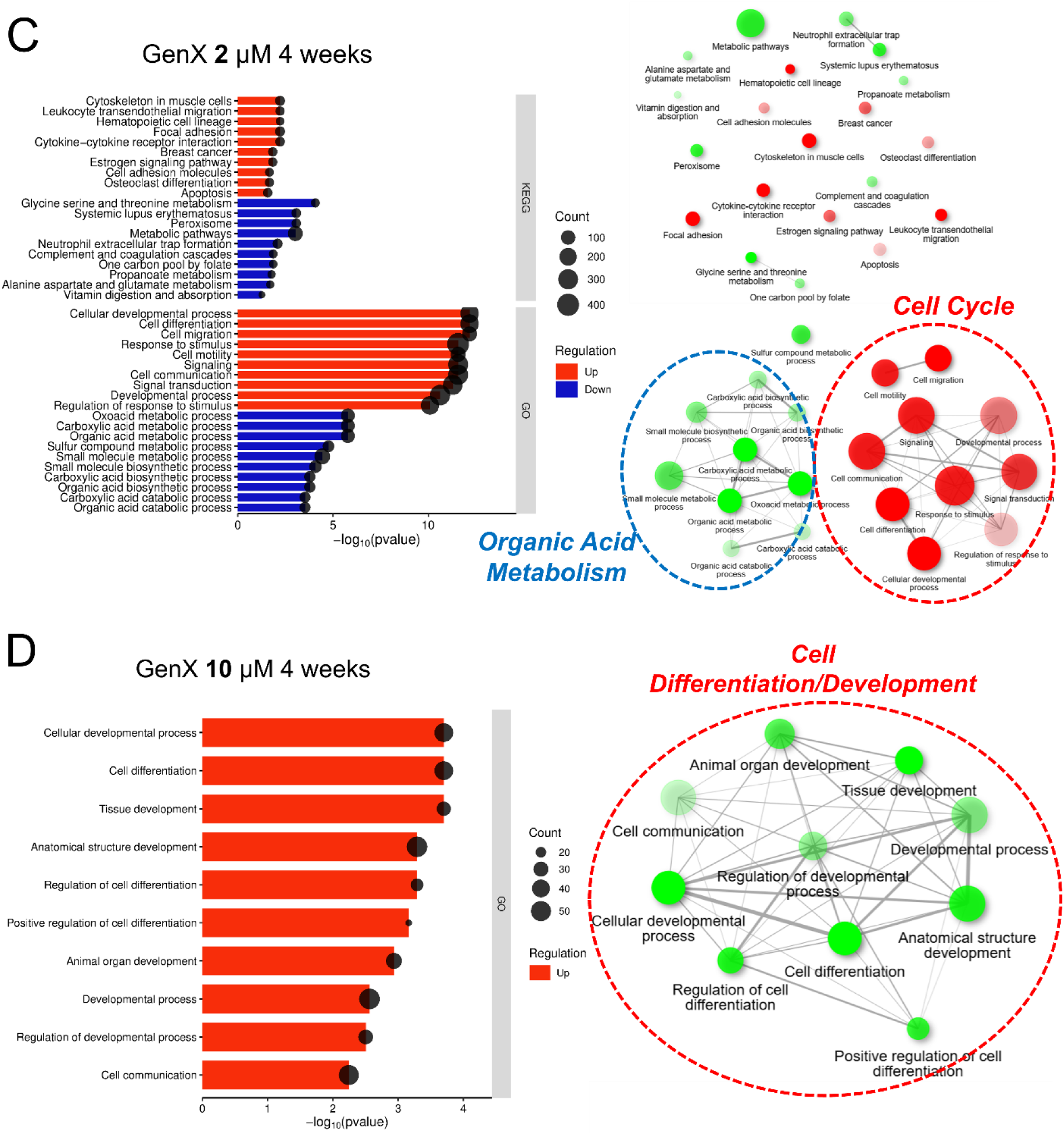
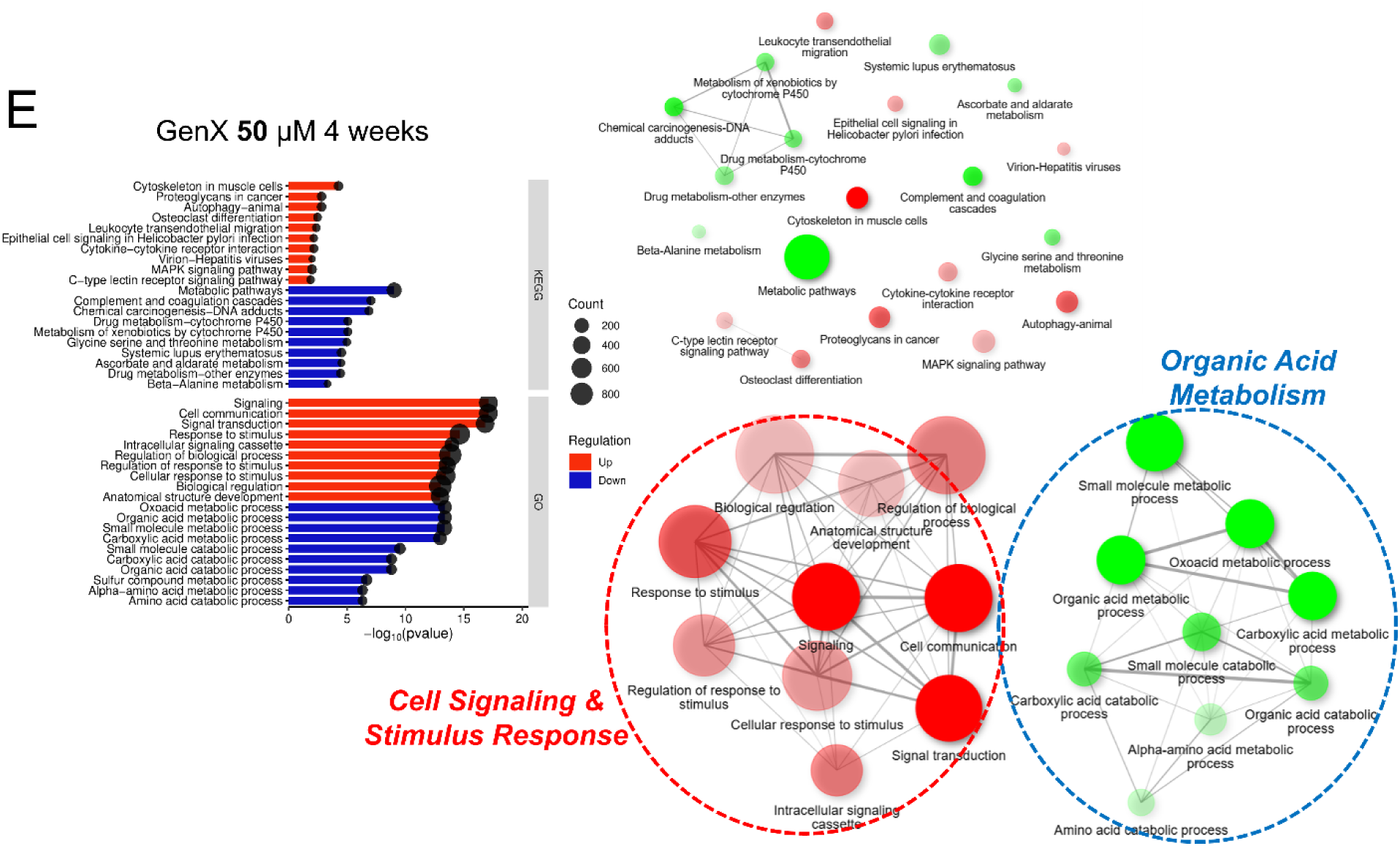
Chronic GenX exposure progressively shifts biological responses toward metabolic reprogramming and cellular signaling. Functional enrichment analysis of differentially expressed genes following 4 weeks of GenX exposure at 0.08 (A), 0.4 (B), 2 (C), 10 (D), and 50 (E) μM, respectively. Representative enriched KEGG and Gene Ontology Biological Process (GO-BP) terms are shown for each concentration with graphical presentation of clustered KEGG pathways and GO-BP. For the lowest concentration (0.08 μM), additional GO-BP enrichment analysis generated using STRING-DB is also presented. Enriched pathways were identified using DEGs (|log2FC| > 1, FDR < 0.05). Only the top 10 enriched pathways are shown.

At 0.08 µM, enrichment analysis revealed mRNA changes consistent with suppression of lipid metabolism-related pathways, including steroid hormone biosynthesis and glycosphingolipid biosynthesis, together with changes in amino acid metabolism, suggesting that prolonged exposure to environmentally relevant GenX concentrations was sufficient to perturb hepatic metabolic homeostasis (Figure 5A). At 0.4 µM, metabolic pathway remodeling became more pronounced, with additional enrichment of fatty acid metabolism and xenobiotic-related pathways, indicating expansion of the metabolic response beyond the lowest exposure concentration (Figure 5B). At the intermediate concentration (2 µM), broader perturbation of lipid metabolism and cellular metabolic processes was observed, accompanied by increasing enrichment of pathways involved in cellular signaling and homeostatic regulation (Figure 5C).

Although relatively fewer significantly enriched pathways were detected at 10 µM, similar to the acute response, extensive transcriptional alterations remained evident, suggesting transient attenuation of coordinated pathway enrichment despite substantial gene expression changes (Figure 5D). At the highest concentration (50 µM), chronic GenX exposure induced widespread suppression of metabolic pathways, including cytochrome P450-mediated xenobiotic metabolism, amino acid metabolism, and lipid metabolic processes, together with enrichment of multiple signaling pathways associated with cellular stress and structural remodeling (Figure 5E). Overall, chronic GenX exposure produced a coordinated concentration-dependent transcriptional reprogramming characterized by an early disruption of metabolic homeostasis at environmentally relevant concentrations followed by broad metabolic suppression and activation of cellular signaling pathways at higher concentrations.

### 3.5. Exposure Duration Drives Progressive Transcriptomic Divergence Following GenX exposure

To evaluate whether acute transcriptional responses could predict long-term outcomes, we performed correlation analyses comparing acute (4 days) and chronic (4 weeks) transcriptomic profiles across matched exposure concentrations (Supplementary Figure S2). We first evaluated correlations among biological replicates within each exposure condition and demonstrated excellent reproducibility, with Pearson correlation coefficients exceeding 0.99 for both control (0 μM) and high-dose (50 μM) groups (Figure S2A–S2B and Figure S2D–S2E). To assess the intrinsic stability of the steady state mRNA expression levels of the 3D liver spheroid model over time, untreated control samples collected after 4 days and 4 weeks were compared. Control spheroids maintained a highly stable transcriptional profile throughout the culture period, exhibiting a strong correlation (R = 0.98) between time points (Figure S2C). These findings indicate that prolonged culture alone had minimal impact on global gene expression patterns. In contrast, GenX exposure introduced a progressive temporal divergence that became increasingly apparent at higher concentrations. At 50 μM, the correlation between acute and chronic transcriptomic profiles decreased relative to untreated controls (R = 0.95), accompanied by increased dispersion of individual genes from the diagonal distribution (Figure S2D and S2F). Although the overall correlation remained high, the increased scatter indicates that prolonged exposure resulted in substantial mRNA changes beyond the initial acute response.

These findings demonstrate that chronic GenX exposure does not simply amplify acute mRNA changes but instead induces a distinct mRNA expression profile that emerges over time. Together with the differential pathway enrichment and DEG analyses, the correlation results identify exposure duration as an independent determinant of GenX-induced transcriptomic changes and support the importance of incorporating chronic exposure models into GenX hazard assessment.

### 3.6. Benchmark dose modeling reveals time-dependent expansion of transcriptional responsiveness

To evaluate the influence of exposure duration on GenX-induced transcriptional responses, benchmark dose (BMD) distributions were compared between acute (4 days) and chronic (4 weeks) exposures using cumulative accumulation plots generated by BMDExpress 3.0 (Figure 6). Chronic exposure produced a marked upward shift in the accumulation curves relative to the acute condition, indicating that a substantially greater number of genes met benchmark dose modeling criteria. Differences were also evident in the distribution of BMD estimates. The chronic exposure curves were shifted toward lower BMD, benchmark dose lower confidence limit (BMDL) (Supplementary Figure S3A), and benchmark dose upper confidence limit (BMDU) (Supplementary Figure S3B) values compared with the acute exposure, indicating that a larger proportion of transcriptional responses occurred within lower dose ranges. In addition, the chronic condition exhibited a steeper accumulation pattern across intermediate BMD values, reflecting a greater concentration of responsive genes within a relatively narrow benchmark dose range.

**Figure 6.**
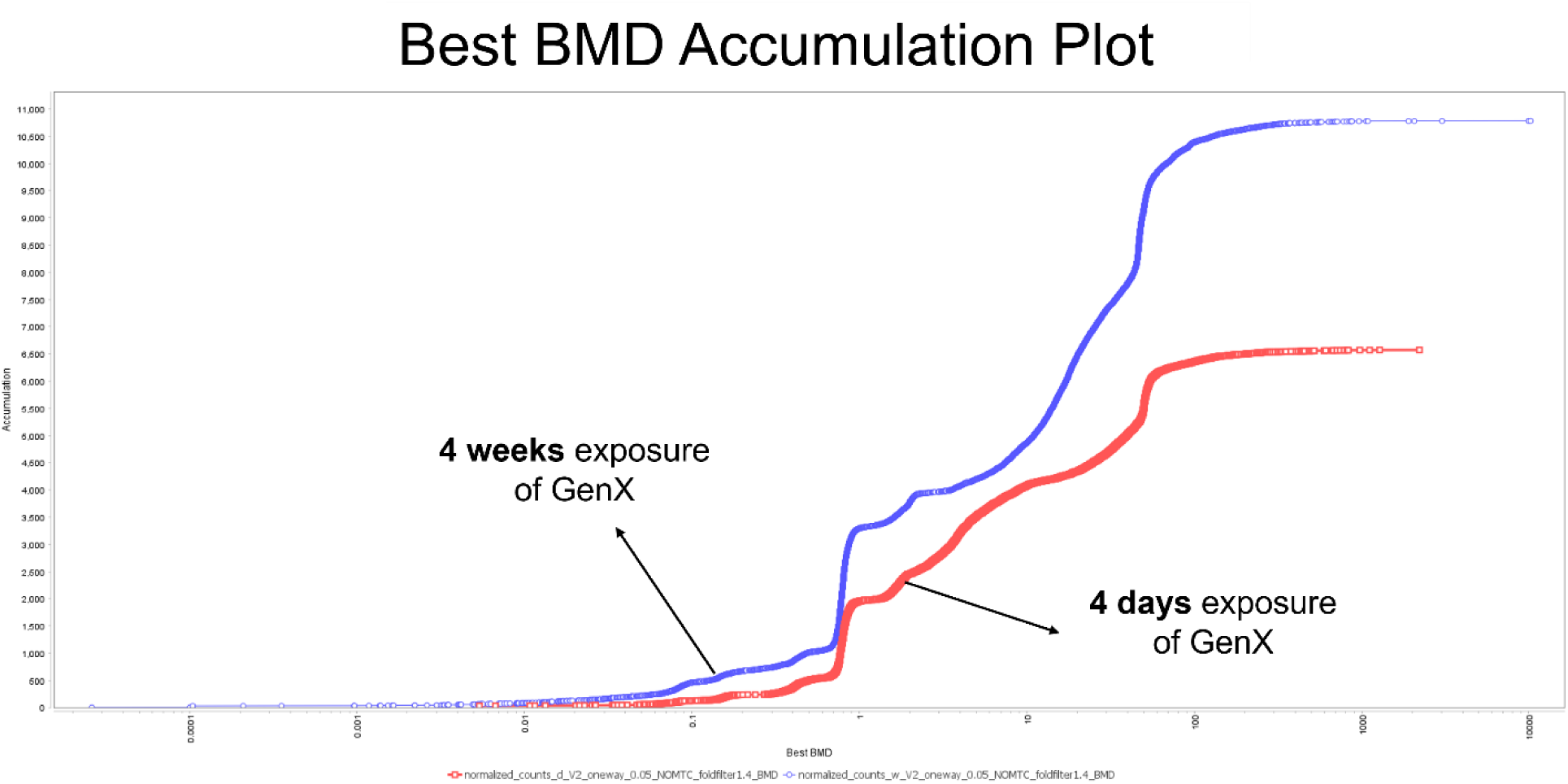
Benchmark dose (BMD) accumulation analyses for acute and chronic GenX exposure. Best benchmark dose (BMD) generated from transcriptomic benchmark dose modeling of GenX-exposed 3D human liver spheroids. Curves represent the cumulative number of genes meeting benchmark dose modeling criteria across increasing BMD values. Red lines indicate acute exposure (4 days), and blue lines indicate chronic exposure (4 weeks). Benchmark dose modeling was performed using normalized gene expression data and the best-fitting model selected for each gene according to BMDExpress criteria^46^.

In contrast, acute exposure produced a more gradual accumulation profile and a smaller overall number of modeled genes (i.e., the number of accumulated genes presented in y-axis of Figure 6). The higher plateau observed following chronic exposure further demonstrates a broader transcriptional response after prolonged GenX treatment. Collectively, these findings indicate that exposure duration substantially influences the distribution and extent of GenX-induced transcriptional responses, with chronic exposure resulting in a larger number of benchmark dose-responsive genes and a distinct BMD distribution compared with acute exposure.

## 4. Discussion

The present study provides systematic comparisons of acute and chronic GenX exposure in a human-relevant 3D liver spheroid model across environmentally relevant and mechanistically informative concentrations. Three principal findings emerged from our analysis. First, exposure duration was a major determinant of transcriptomic outcome, with acute and chronic exposure producing qualitatively distinct molecular responses. Second, GenX elicited non-monotonic transcriptomic responses during acute exposure, whereas chronic exposure generated broader and more coordinated dose-dependent effects. Third, prolonged exposure increased transcriptomic sensitivity and revealed mRNA changes consistent with lipid metabolic perturbations and pathway-level responses that were not apparent during short-term exposure. Collectively, these findings demonstrate that both concentration and exposure duration are likely critical determinants of GenX associated transcriptomic responses and these factors should be considered in the application of transcriptomic based indicators from NAMs in human health risk assessment.

One of the most notable findings was the pronounced non-monotonic response observed following acute exposure. The largest transcriptional perturbation occurred at 0.4 μM rather than at the highest tested concentration, indicating that biological responses did not increase proportionally with dose. Similar non-monotonic behavior has increasingly been reported for PFAS and other environmental contaminants, presenting a significant challenge for toxicological risk assessment because low- and intermediate-dose responses cannot necessarily be predicted from high-dose studies alone^57–61^. Acute exposure also exhibited clear concentration-specific biological responses. Low concentrations primarily result in mRNA changes in proteins involved in cell cycle-associated pathways, suggesting rapid adaptive and proliferative signaling, whereas intermediate concentrations preferentially altered lipid metabolic pathways, including fatty acid metabolism and PPAR signaling. Additional STRING-DB GO-BP network analysis further demonstrated that these low-dose responses in acute exposure were centered on interconnected biological processes involved in mitotic cell cycle regulation, chromosome organization, and cell division, suggesting that environmentally relevant GenX concentrations perturb proliferative regulatory networks rather than overt stress-response pathways. At the highest concentration, stress- and toxicity-associated pathways such as PI3K-Akt, ErbB, and Toll-like receptor signaling became dominant. PI3K-Akt signaling pathway has been suggested as one of potential PFAS-induced toxicity mechanisms, which also regulates apoptosis and cellular proliferation^62,63^. This progression suggests a transition from early adaptive responses to metabolic disruption and ultimately activation of cellular stress-survival programs. The non-monotonic nature of these responses further implies that distinct biological programs may be engaged across different exposure levels rather than representing a simple increase in response magnitude.

In contrast, chronic GenX exposure elicited broader and more coordinated transcriptional responses characterized by early and persistent perturbation of lipid metabolic pathways. Even the lowest concentrations altered mRNA expression of proteins in pathways associated with glycosphingolipid metabolism, steroid metabolism, fatty acid metabolism, and peroxisomal function, supporting the growing consensus that disruption of lipid homeostasis represents a central mechanism of PFAS toxicity^64,65^. Unlike the concentration-specific patterns observed during acute exposure, chronic exposure produced a largely dose-dependent response profile, consistent with progressive cellular adaptation and cumulative biological effects. Notably, low-dose chronic exposure primarily affected pathways associated with metabolic homeostasis and lipid regulation, whereas higher concentrations activated stress-response and toxicity-related pathways with minimal overlap between the two response domains. These findings suggest a mechanistic transition from adaptive metabolic remodeling to overt cellular stress and dysfunction as exposure levels increase in a similar pattern with the acute exposure condition. Furthermore, prolonged exposure was characterized by suppression of diverse metabolic pathways accompanied by activation of inflammatory and stress-response signaling, indicative of a progressive loss of hepatocellular metabolic identity^66,67^. Similar metabolic reprogramming and fibroinflammatory signatures have been reported in primary human hepatocytes exposed to GenX^20^, supporting the biological relevance of the transcriptional responses observed in the present spheroid model. Notably, the presence of transcripts that were consistently altered across all exposure concentrations suggests that a subset of cellular responses to GenX is highly robust and largely independent of dose magnitude, potentially representing core molecular events associated with GenX exposure rather than secondary concentration-dependent effects. Moreover, the dramatic increase in the number of constitutively responsive transcripts during chronic exposure relative to acute exposure suggests convergence towards a stable mRNA expression level changes that become progressively independent of dose magnitude. This finding supports the notion that prolonged GenX exposure promotes sustained biological reprogramming rather than transient adaptive responses. For example, the sustained increased of expression level of circadian rhythm genes following chronic GenX exposure is notable because circadian regulators are closely linked to hepatic metabolism, xenobiotic responses, and lipid homeostasis^68,69^, suggesting that dysregulation of clock-associated transcriptional programs may contribute to the metabolic reprogramming observed following prolonged exposure.

Rather than representing a simple extension of the acute response, chronic GenX exposure shifted the transcriptomic landscape from early adaptive responses toward a coordinated metabolic reprogramming that became evident even at environmentally relevant concentrations. Functional enrichment analysis further demonstrated that exposure duration substantially altered the biological processes affected by GenX. During acute exposure, low concentrations preferentially cell cycle-associated pathways, suggesting an adaptive proliferative response, whereas intermediate concentrations increasingly perturbed lipid metabolic pathways before shifting toward endocrine- and signaling-related processes at the highest concentration. In contrast, chronic exposure produced a progressive transition toward coordinated metabolic reprogramming characterized by enrichment of organic acid metabolic processes, developmental programs, and cell signaling pathways. Because fatty acids represent a major class of organic acids, enrichment of organic acid metabolism is consistent with increasing evidence that chronic PFAS exposure disrupts hepatic lipid homeostasis and broader intermediary metabolism. The emergence of cell signaling and stimulus-response pathways after prolonged exposure further suggests that chronic GenX exposure induces coordinated regulatory responses beyond simple dose-dependent toxicity, supporting the concept that sustained low-dose exposure engages biological programs that are not fully captured in conventional short-term assays.

Perhaps the most important finding from a risk assessment perspective was the limited concordance between acute and chronic responses. Many pathways enriched after 4 weeks of exposure were absent or only weakly represented during acute exposure, and transcriptomic correlation analyses demonstrated increasing divergence between acute and chronic transcriptional states with increasing GenX concentration. These results suggest that chronic toxicity cannot be reliably inferred from short-term studies and instead reflects secondary and adaptive molecular programs that emerge during sustained exposure. Therefore, these findings suggest that exposure duration should be considered not only when interpreting transcriptomic mechanisms but also when applying tPODs in chemical risk assessment, as short-term transcriptomic responses may not fully reflect the coordinated metabolic reprogramming associated with chronic exposure. Our results also support the growing recognition that chronic low-dose PFAS exposure may engage unique biological processes that cannot be predicted from acute or high-dose studies alone^70,71^. Consistent with this interpretation, benchmark dose modeling (BMD) demonstrated that chronic exposure lowered transcriptomic points of departure and expanded the number of affected biological pathways, indicating that prolonged exposure increased transcriptomic sensitivity to GenX. That is, Chronic exposure resulted in a leftward shift of transcriptomic BMD distributions, suggesting increased transcriptional sensitivity and expanded biological engagement following prolonged GenX exposure. As transcriptomic points of departure become increasingly incorporated into next-generation risk assessment (NRGA) frameworks^72^, our findings suggest that tPODs derived from chronic human-relevant spheroid models may provide more biologically informative and health-protective estimates of chemical potency than those generated from conventional acute exposure systems. More broadly, the data indicates that prolonged GenX exposure unmasks molecular responses associated with metabolic dysfunction and loss of hepatocellular identity that are not evident during short-term exposure, highlighting the importance of incorporating exposure duration into PFAS hazard characterization and human health risk assessment.

## Data availability

Raw and processed RNA-seq data are currently being deposited in the NCBI Gene Expression Omnibus (GEO) database under the accession number, GSE339046.

